# A copy number variant is associated with a spectrum of pigmentation patterns in the rock pigeon (*Columba livia*)

**DOI:** 10.1101/688671

**Authors:** Rebecca Bruders, Hannah Van Hollebeke, E.J. Osborne, Zev Kronenberg, Mark Yandell, Michael D. Shapiro

**Affiliations:** School of Biological Sciences, University of Utah, Salt Lake City, Utah, United States of America; Department of Human Genetics, University of Utah, Salt Lake City, Utah, United States of America

## Abstract

Rock pigeons (*Columba livia*) display an extraordinary array of pigment pattern variation. One such pattern, Almond, is characterized by a variegated patchwork of plumage colors that are distributed in an apparently random manner. Almond is a sex-linked, semi-dominant trait controlled by the classical *Stipper* (*St*) locus. Heterozygous males (Z*^St^*Z*^+^* sex chromosomes) and hemizygous Almond females (Z*^St^*W) are favored by breeders for their attractive plumage. In contrast, homozygous Almond males (Z*^St^*Z*^St^*) develop severe eye defects and lack all plumage pigmentation, suggesting that higher dosage of the mutant allele is deleterious. To determine the molecular basis of Almond, we compared the genomes of Almond pigeons to non-Almond pigeons and identified a candidate *St* locus on the Z chromosome. We found a copy number variant (CNV) within the differentiated region that captures complete or partial coding sequences of four genes, including the melanosome maturation gene *Mlana*. We did not find fixed coding changes in genes within the CNV, but all genes are misexpressed in regenerating feather bud collar cells of Almond birds. Notably, six other alleles at the *St* locus are associated with depigmentation phenotypes, and all exhibit expansion of the same CNV. Structural variation at *St* is linked to diversity in plumage pigmentation and gene expression, and thus provides a potential mode of rapid phenotypic evolution in pigeons.

**AUTHOR SUMMARY:** The genetic changes responsible for different animal color patterns are poorly understood, due in part to a paucity of research organisms that are both genetically tractable and phenotypically diverse. Domestic pigeons (*Columba livia*) have been artificially selected for many traits, including an enormous variety of color patterns that are variable both within and among different breeds of this single species. We investigated the genetic basis of a sex-linked color pattern in pigeons called Almond that is characterized by a sprinkled pattern of plumage pigmentation. Pigeons with one copy of the Almond allele have desirable color pattern; however, male pigeons with two copies of the Almond mutation have severely depleted pigmentation and congenital eye defects. By comparing the genomes of Almond and non-Almond pigeons, we discovered that Almond pigeons have extra copies of a chromosome region that contains a gene that is critical for the formation of pigment granules. We also found that different numbers of copies of this region are associated with varying degrees of pigment reduction. The Almond phenotype in pigeons bears a remarkable resemblance to Merle coat color mutants in dogs, and our new results from pigeons suggest that similar genetic mechanisms underlie these traits in both species. Our work highlights the role of gene copy number variation as a potential driver of rapid phenotypic evolution.

## INTRODUCTION

In natural populations of animals, pigment colors and patterns impact mate choice, signaling, mimicry, crypsis, and distraction of predators [1, 2]. In domestic animals, pigmentation traits are often selected by humans based on colors and patterns they find most attractive. Despite longstanding interest in the spectactular variation in color and pattern among animals, we know little about the molecular mechanisms that mediate color patterns. Understanding the genetic basis of the stunning array of animal color patterns benefits from the study of genetically tractable species; however, progress is hampered, in part, by a limited number of traditional model organisms that show limited variation in color and color patterning.

The domestic rock pigeon (*Columba livia*) is a striking example of variation shaped by artificial selection, with a multitude of colors and color patterns within and among more than 350 breeds. Because breeds of domestic pigeon belong to the same species and are interfertile, pigeons offer an exceptional opportunity to understand the genetic basis of pigmentation traits using laboratory crosses and genomic association studies [3]. Previously, we identified several genes involved in determining the type and intensity of plumage melanins in pigeons [4, 5], but considerably less is known about the molecular determinants of pattern deposition [6]. The molecular basis of pattern variation is an exciting frontier in pigmentation genetics, and recent work in other vertebrates reveals several genes that contribute to this process. Still, the genetic basis of pigment pattern is decidedly less well understood than the genes controlling pigment types [7–16].

The classical pigmentation pattern in *C. livia* known as Almond is caused by a semi-dominant mutation (*St* allele) at the sex-linked *Stipper* (*St*) locus [17] (Fig. 1). Unlike most other pigmentation pattern traits in pigeons, the variegated or sprinkled patchwork of plumage colors in Almond is apparently random within and among individuals [18]. Furthermore, the color pattern changes in an unpredictable manner with each molt [19–21]. The number of pigmented feathers in Almond pigeons also increases with each successive molt, and this effect is more pronounced in males [22, 23]. Notably, this phenomenon is the opposite of what is typically observed with pigmentation traits that change throughout the lifespan of an individual, such as vitiligo and graying, which result in a decrease in pigment over time [24–28]. In addition to Almond, at least six other alleles at *St* lead to varying degrees of depigmentation in pigeons, suggesting that the *St* locus might be a mutational hotspot [21, 29].

**Figure 1.**
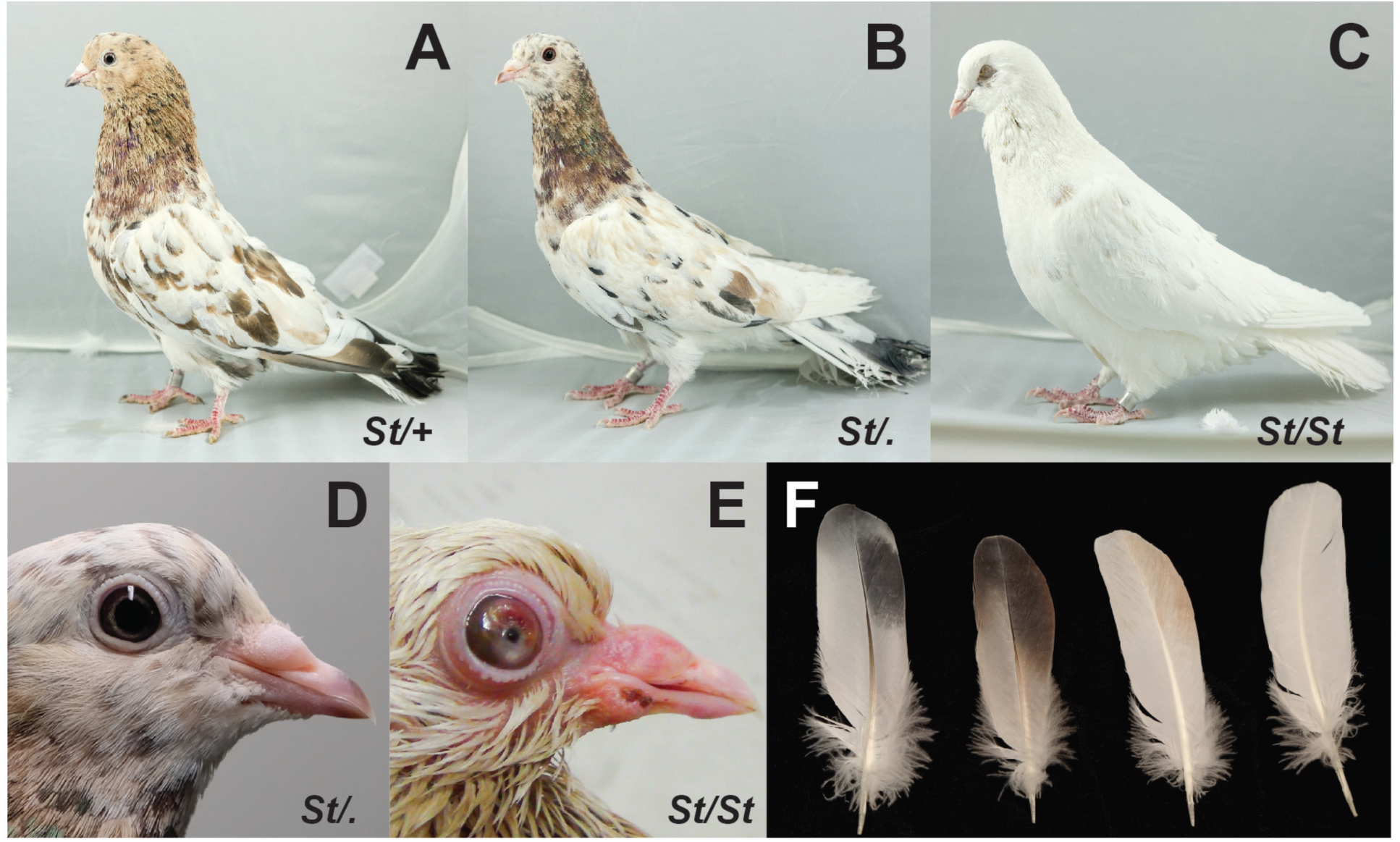
Phenotypes of pigeons carrying Almond alleles (*St*, Almond allele; +, wild type allele). (A) Heterozygous Almond male. (B) Hemizygous Almond female. (C) Homozygous Almond male. (D) Almond females have no observable eye defects. (E) Homozygous Almond males often show severe eye defects. Defects pictured in this juvenile include bloated eyelid and anterior opacity. (F) Wing feathers from different phenotypes, left to right: non-Almond, dark Almond, light Almond, homozygous Almond.

Heterozygous Almond males (Z*^St^*Z*^+^*) and hemizygous Almond females (Z*^St^*W; males are the homogametic sex in birds), each of which have one copy of the *St* allele, are valued by breeders for their attractive color patterns. However, homozygous Almond males (Z*^St^*Z*^St^*) almost always lack pigmentation in the first set of pennaceous feathers and have severe congenital eye defects [19, 30, 31] (Fig. 1B, C). The pattern of inheritance of Almond suggests that dosage of the mutant allele, rather than absence of the wild type allele, is responsible for the pigment and eye phenotypes in homozygous males. Eye defects are also associated with pigmentation traits in other vertebrate species, including dogs and horses, yet the molecular basis of these linked effects remains poorly understood [9, 32–37]. Therefore, Almond pigeons can illuminate links between pigmentation and eye defects, including whether pleiotropic effects of a single gene or linked genes with separate effects control these correlated traits.

In this study, we investigate the genomic identity of the *St* locus in domestic pigeons. Whole-genome sequence comparisons of Almond and non-Almond birds reveal a copy number variant (CNV) in Almond birds that includes the complete coding sequences of two genes, and partial coding sequences of two others. One of the complete genes, *Mlana*, plays a key role in the development of the melanosome (the organelle in which pigment granules are produced), making it a strong candidate for the pigmentation phenotype observed in Almond pigeons. We also find that different alleles at *St* are correlated with different degrees of expansion of the same CNV, thereby linking a spectrum of pigmentation variants to changes at one locus.

## RESULTS

### A sex-linked genomic region is associated with Almond pigmentation pattern

To determine the genomic location of the sex-linked *St* locus, we compared the genomes of 12 Almond pigeons to a panel of 109 non-Almond pigeons from a diverse set of breeds, using a probabilistic measure of allele frequency differentiation (pFst) [38] (see S1 Table for sample details). This whole-genome scan identified several significantly differentiated regions, but one exceeded the others by several orders of magnitude and was located on a Z-chromosome scaffold (ScoHet5_227), as predicted from classical genetics studies (Fig. 2A). The differentiated region of ScoHet5_227 (position 5,186,219-5,545,482; peak SNP, *p*=1.1 e-16, genome wide significance threshold p = 5.5 e-10) contained eight annotated protein-coding genes, none of which had fixed coding changes in Almond compared to non-Almond genomes (VAAST [39]). Therefore, the Almond pigmentation pattern probably does not result from non-synonymous changes to protein-coding genes.

**Figure 2.**
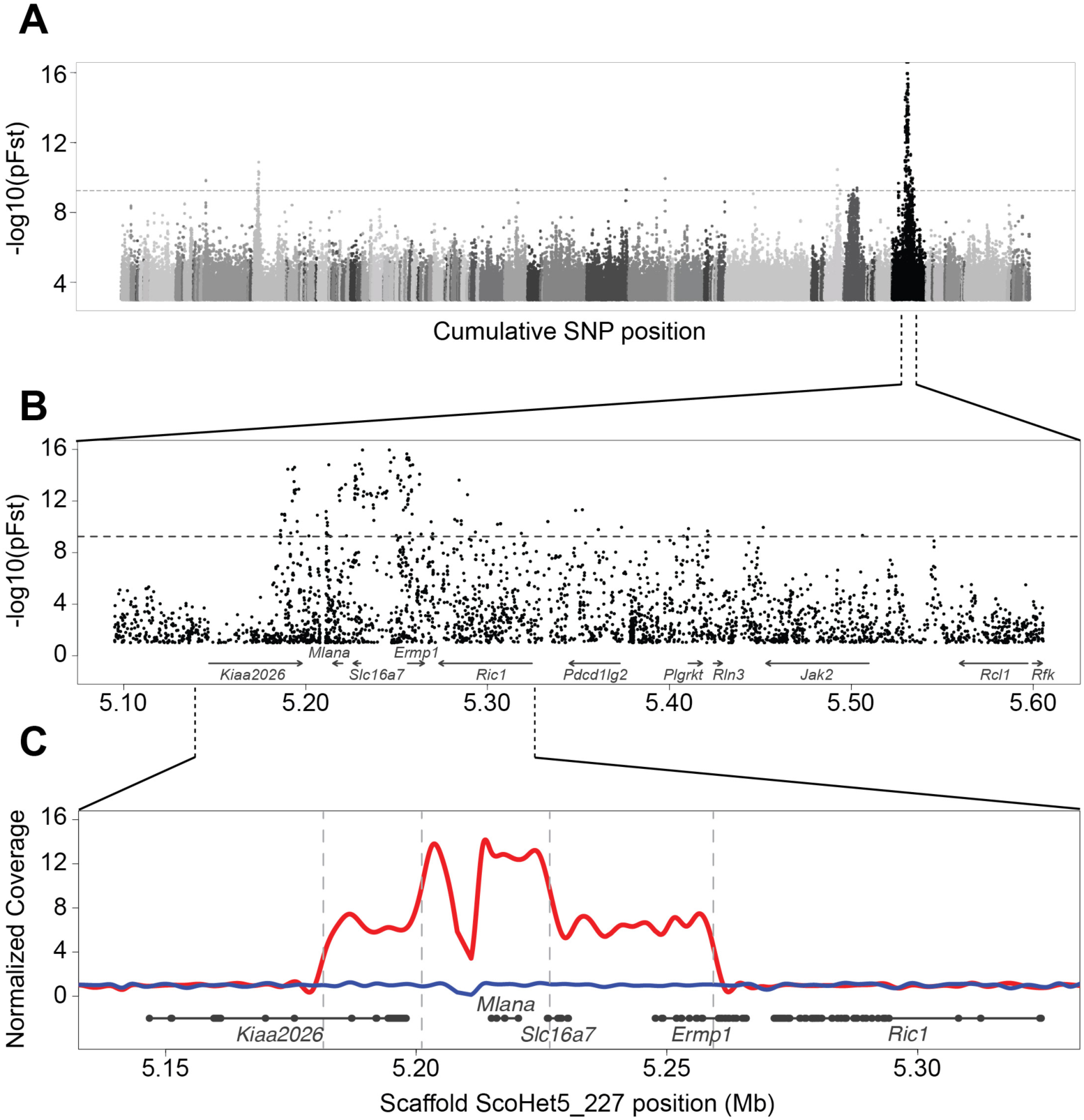
Almond is associated with a CNV on a sex-linked genomic scaffold. (A) Whole-genome pFst comparisons between Almond and non-Almond pigeons. Each dot represents a SNP position, with shades of gray indicating different genomic scaffolds. The horizontal dashed grey line indicates genome-wide significance threshold. (B) Detail of pFst plot for candidate region on ScoHet5_227, a sex-linked scaffold [104]. Gene models are depicted at the bottom of the plot. (C) Detail view of the CNV region. Solid red line represents the mean normalized read depth for 10 female Almond birds in this region. The blue line is a single representative of non-Almond female coverage. Vertical dashed lines indicate positions of CNV breakpoints. Gene models are depicted below the coverage plot in grey (thick lines, exons; thin lines, introns).

Several other scaffolds contained sequences that were significantly differentiated between Almond and non-Almond pigeons (Fig. 2A). All of these regions are autosomal, and we speculate that they are linked to other color traits that are often co-selected with Almond to give the most desirable sprinkled patchwork of colors, including T-check (a highly melanistic wing pattern), kite bronze (a deep reddening of the feathers), and recessive red (a pheomelanic color trait) [18, 21, 29]. However, because Almond is a sex-linked trait [17], we focused our attention on the Z-linked scaffold ScoHet5_227.

### A copy number variant is associated with the Almond pigment pattern

In the absence of fixed coding changes between Almond and non-Almond birds, we next asked if birds with different phenotypes had genomic structural differences in the candidate region. We examined sequencing coverage on ScoHet5_227 and found that all 12 Almond genomes had substantially higher coverage in the Almond candidate region relative to non-Almond genomes, indicating the presence of a copy number variant (CNV) (Fig. 2B, C). The CNV captures a 77-kb segment of the reference genome (ScoHet5_227: 5,181,467-5,259,256), with an additional increase in coverage in a nested 25-kb segment (ScoHet5_227: 5,201,091-5,226,635). Read-depth analysis confirmed 7 copies of the outer 77-kb segment and 14 copies of the inner 25-kb segment in the genomes of female (Z*^St^*W) Almond pigeons, which have an *St* locus on only one chromosome. We used PCR to amplify across the outer and inner CNV breakpoints of Almond pigeons and determined that the CNV consists of tandem repeats of the 77-kb and nested 25-kb segments (Fig. 3B).

**Figure 3.**
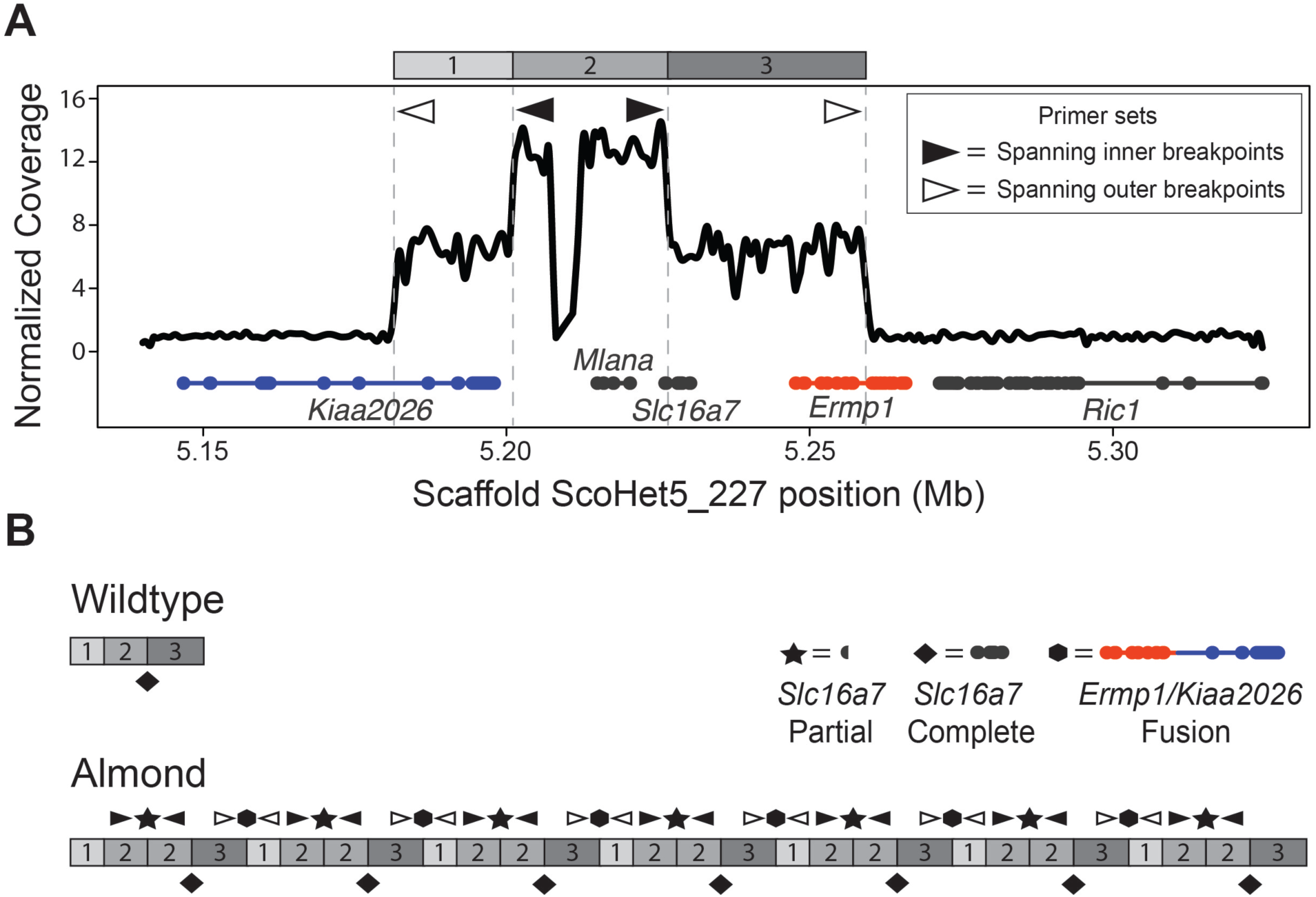
The Almond-associated CNV has a complex structure that results in duplicated, truncated, and fused genes. (A) Coverage diagram showing different regions of the CNV normalized to a non-CNV region on the same scaffold. Two outer regions (1 and 3, above plot) have an approximately 7-fold coverage increase, while one inner region (2) has an approximately 14-fold coverage increase. Gene models are depicted below the coverage plot in grey, orange and blue (thick lines, exons; thin lines, introns). (B) Schematic of the non-Almond (top) and inferred Almond (bottom) structures of the CNV. Gene structural changes resulting from the Almond CNV include a fusion of *Ermp1* and *Kiaa2026* at the segment 3/1 junction (hexagon), and a truncated version of *Slc16a7* at the segment 2/2 junction (star). A complete copy of *Slc16a7* occurs at each 2/3 junction (diamond).

We then genotyped the CNV region in a larger sample of Almond pigeons and found a significant association between the number of tandem repeats and the Almond phenotype (TaqMan assay; pairwise Wilcoxon test, *p*= 2.0 e-16). Almost all Almond birds have more than one copy of the CNV per Z-chromosome (n=78 of 80) (Fig. 4). Conversely, nearly all non-Almond birds had only one copy per Z-chromosome (n=55 of 57). The two non-Almond birds with >1 copy per chromosome had a maximum of one additional copy of the CNV, indicating that small increases in copy number do not necessarily cause the Almond phenotype. Overall, these analyses suggest that expansion of CNV on ScoHet5_227 is associated with the Almond phenotype.

**Figure 4.**
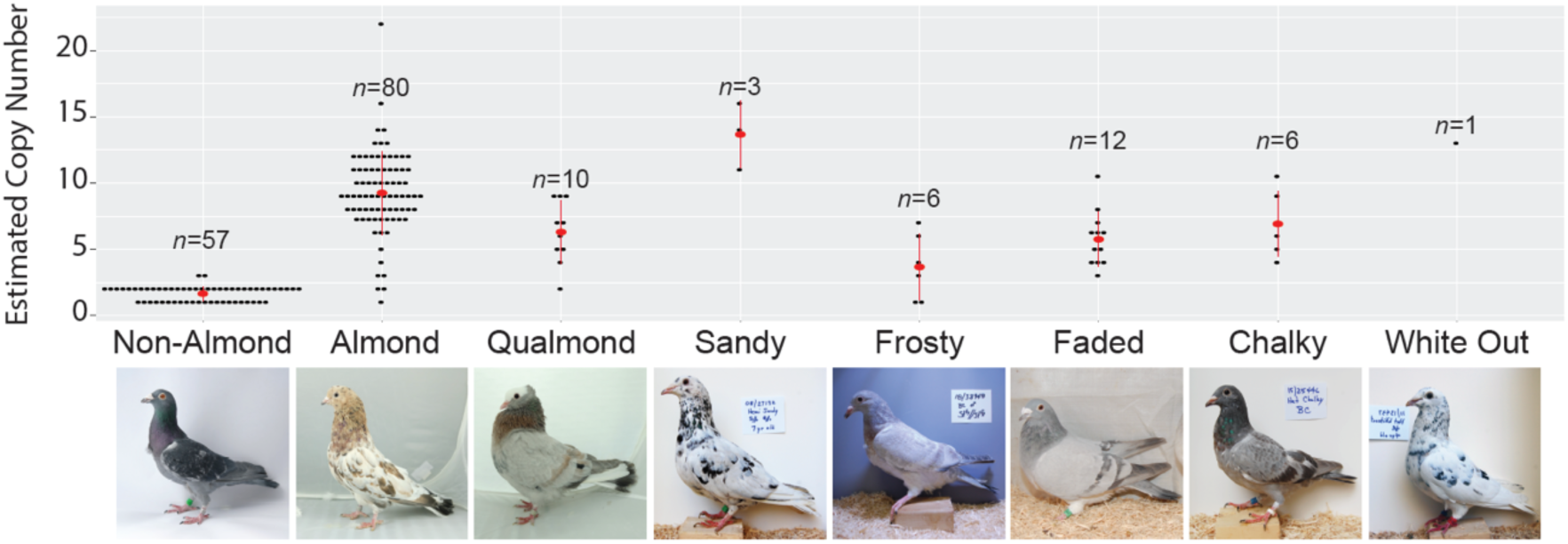
*St*-linked pigmentation phenotypes show quantitative variation in the Almond CNV. Black dots represent results of a TaqMan copy number assay. Mean copy numbers for each phenotype are shown as red dots. Most individuals without *St*-linked phenotypes have the expected 1 or 2 copies (because *St* is a sex-linked locus, females have a minimum of 1 copy and males have a minimum of 2). All other *St*-linked phenotypes are associated with an expansion of the CNV in the Almond candidate region on scaffold ScoHet5_227, indicating an allelic series at *St*. Numbers above each phenotype indicate number of individuals sampled.

### Genes within the CNV are misexpressed in Almond feather buds

We next asked if the CNV was associated with gene expression changes between developing Almond and non-Almond feathers. To address this question, we compared expression of genes in the CNV region among birds with (Z*^St^*Z*^+^*, Z*^St^*W, Z*^St^*Z*^St^*) and without (Z*^+^*Z*^+^*and Z*^+^*W) Almond alleles. We analyzed Almond feather buds with dark and light pigmentation separately to assess whether expression differed between qualitatively different feather pigmentation types, both of which are present in Z*^St^*Z*^+^*and Z*^St^*W Almond individuals. The CNV contains the complete coding sequences of two genes, *Mlana* and *Slc16a7*, and partial coding sequences of two additional genes, *Ermp1* and *Kiaa2026* (Fig. 3A). *Mlana* is predicted to have up to 14 total copies per Z*^St^* chromosome based on sequencing coverage in Z*^St^*W Almond birds (Fig. 3B). *Mlana* is expressed almost exclusively in melanocytes (melanin-producing cells), and encodes a protein that is critical for melanosome maturation through interactions with the matrix-forming protein Pmel [40–42]. Thus, the combination of the biological role of *Mlana* and its location in the Almond CNV makes *Mlana* a strong candidate gene for the Almond phenotype.

Compared to non-Almond feather buds, *Mlana* expression is increased in dark feather buds, but not in light feather buds, from Z*^St^*Z*^+^* and Z*^St^*W Almond birds or the unpigmented feather buds of homozygous Almond (Z*^St^*Z*^St^*) birds (Fig. 5A; see S2 Table and S3 Table for raw data for all qRT-PCR experiments). We noticed that the variance of expression observed for *Mlana* in both dark and light Almond feather buds, though not statistically significant (Kolmogorov-Smirnov test), trends higher than in non-Almond samples. This data distribution might reflect the variability of the phenotype itself, which is characterized by different quantities and intensities of feather pigmentation both within and between Z*^St^*Z*^+^*and Z*^St^*W Almond pigeons.

**Figure 5.**
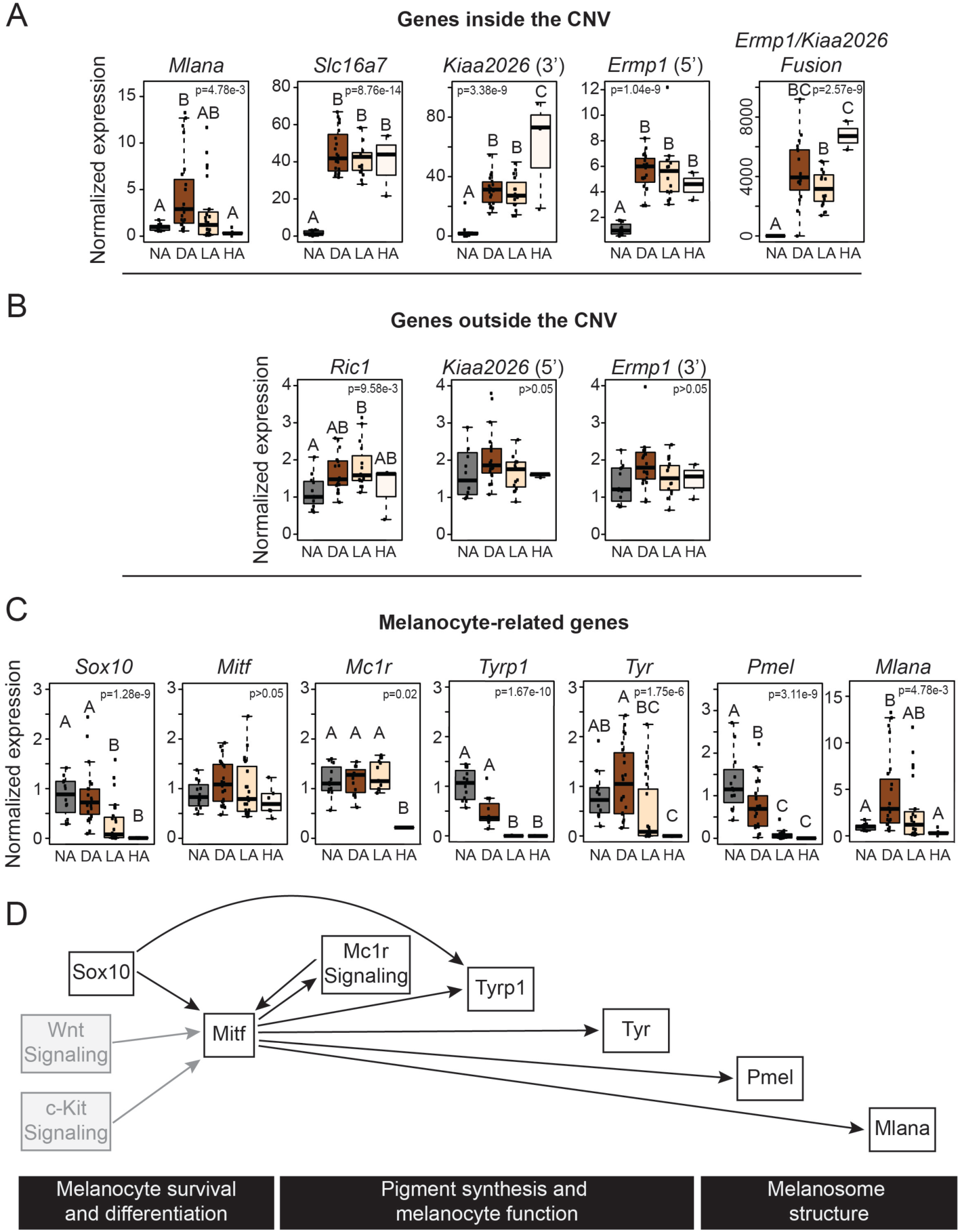
Almond and non-Almond feather buds have distinct gene expression profiles. (A) Exons assayed within the CNV show expression differences in Almond feather buds compared to non-Almond. Boxplots show the results of qRT-PCR assays designed to assess gene expression of exons located in the CNV region. Fusion gene expression results are from qPCR primers spanning exon 7 of *Ermp1* into exon 5 of *Kiaa2026.* (B) Exons assayed outside the CNV show no expression differences in Almond feather buds compared to non-Almond. This indicates expression differences are specific to exons inside the CNV. (C) Expression of melanocyte-related genes. qRT-PCR results indicate a decrease in expression of several genes involved in melanin production in Almond feather buds. (D) Model of interactions among genes and signaling pathways involved in different aspects of pigment synthesis. Gray boxes indicate pathways discussed in the text but not directly represented in our expression analyses. NA, feather buds from non-Almond individuals with wild type alleles at *St*; DA, dark Almond feather buds from hemizygous and heterozygous Almond individuals; LA, light Almond feather buds from hemizygous and heterozygous Almond individuals; HA, feather buds from a homozygous Almond individual. Bar in each box represents the median, box ends indicate upper and lower quartiles, whiskers indicate the highest and lowest value excluding outliers. Different letters indicate groups with statistically significant differences in gene expression determined by ANOVA and post-hoc Tukey test (p<0.05).

Other genes completely or partially within the CNV show increased expression in feathers from birds with at least one Almond allele relative to non-Almond birds. *Slc16a7* encodes a monocarboxylate transporter, and is predicted to be amplified to six full-length copies in Almond pigeons (Fig. 3B). We observed a 40-fold increase in expression of *Slc16a7* in Almond feather buds compared to non-Almond (Fig. 5A). *Slc16a7* is not known to be important in pigmentation; however, this gene is expressed in the mammalian, where it is involved in lactic acid transport and osmotic balance [43–47].

In addition to the two genes fully contained within the CNV, a novel fusion of *Ermp1* (a metallopeptidase gene) and *Kiaa2026* (unknown function) is predicted to span the outer CNV breakpoints (Fig. 3C). Neither gene is known to play a role in pigmentation or eye development. The predicted Ermp1/Kiaa2026 fusion protein is a truncated version of Ermp1, including the peptidase domain and 3 of the 6 transmembrane domains (Fig. 3B). The 22 amino acids from Kiaa2026 at the C-terminus of the fusion protein do not include a known protein domain [48]; thus, the fusion protein is unlikely to create a novel combination of functional domains. As expected, the *Ermp1/Kiaa2026* fusion gene is not expressed in feathers of non-Almond birds, but is expressed in birds with Almond alleles (Fig. 5A). When we analyzed the expression of the exons of *Kiaa2026* and *Ermp1* located outside the CNV, we did not observe expression differences among genotypes (Fig. 5B). Therefore, the Almond CNV is associated with expression of the novel fusion gene, but not with expression differences in the full-length transcripts of either contributing gene. Similarly, *Ric1*, a gene immediately outside the CNV, shows a modest (less than two-fold) expression increase in light Almond feathers relative to other feather types (Fig. 5B). In summary, genes inside CNV show variable or increased expression in feathers from Almond birds, whereas genes adjacent to the CNV show little or no expression change.

### Gene expression changes suggest melanocyte dysfunction in Almond feather buds

Plumage pigmentation patterns in Z*^St^*Z*^+^*, Z*^St^*W, and Z*^St^*Z*^St^* Almond birds are radically different than non-Almond birds, which led us to predict that other components of the melanogenesis pathway might differ as well. The production of melanin by melanocytes is a multi-step process that begins with activation of several pathways, including Wnt and Mc1r signaling, via extracellular ligands and agonists [49–52]. Subsequently, expression of transcription factors, including Mitf, activates a genetic cascade that ultimately promotes the maturation of a functional melanocyte [53]. Within the melanocyte itself, a series of enzymatic reactions and assembly of the melanosome leads to the production and deposition of pigments. Melanosomes are then transferred to skin cells and epidermal appendages, including feathers. In pigeons and other birds with melanin-based pigments, the balance of pheomelanin (reds, yellows) and eumelanin (blacks, browns) deposition determines plumage color [54].

To determine if pigment production signals diverge between Almond and non-Almond feather buds, we measured expression of several marker genes for melanocyte maturation and function by qRT-PCR. We first examined genes involved in melanocyte survival and differentiation, both of which are critical early events in melanin production. *Sox10*, which encodes a transcription factor that activates expression of many downstream genes including *Mitf, Tyrosinase*, and *Tyrp1* expression [55], is downregulated only in light Almond and homozygous Almond feather buds (Fig. 5B). Because *Sox10* regulates *Mitf* and other melanocyte genes, this result indicates that melanocyte dysfunction occurs early in the lightly pigmented Almond feathers, but not in dark Almond feathers. A second melanocyte differentiation and survival marker gene, *Mitf*, encodes a transcription factor that activates expression of *Tyrosinase, Tyrp1, Pmel*, and *Mlana* [40, 53, 56, 57]. Unlike *Sox10*, *Mitf* is not differentially expressed in any of the phenotypes we tested (Fig. 5C). This result suggests that melanocytes are present in the feathers of all phenotypes, even in severely depigmented feathers [9]. Two genes that activate *Mitf* expression (*Sox10* and *Mc1r*, see below; Fig. 5C) are downregulated, which implies that *Mitf* would be downregulated as well. However, the persistence of high *Mitf* expression could be the result of activation by other pathways such as Wnt and c-Kit signaling [56]. Together, our gene expression results indicate that melanocytes are present in all feather buds of Almond pigeons (*Mitf* is expressed), but decreased *Sox10* expression in light and homozygous Almond feathers suggests multiple copies of the Almond CNV are associated with dysfunction early in melanogenesis in light and homozygous Almond feathers (*Sox10* expression is decreased).

We next assayed genes involved in pigment production, an indicator of melanocyte function. *Mc1r*, which encodes a G-protein-coupled receptor necessary for eumelanin production [58], and *Tyrosinase*, which encodes a critical enzyme for both eumelanin and pheomelanin production, were downregulated only in homozygous Almond feather buds (Fig. 5C). Therefore, expression of two key determinants of pigment production is affected only in the most severe depigmentation phenotype. *Tyrp1*, which encodes another enzyme important for eumelanin but not phenomelanin production [59], was downregulated in all Almond feather buds, with the most severe effects in light and homozygous Almond feather buds (Fig. 5C). Thus, the eumelanin synthesis pathway is affected in all Almond feathers, but pigment generation and melanocyte function genes are more impacted in light Almond and homozygous Almond feather buds, with the most severe downregulation observed in homozygotes (Fig. 5C).

Finally, we measured expression of the melanosome structure gene *Pmel*. Melanosome structure is thought to be necessary for eumelanin but not pheomelanin production [60]. Pmel is an amyloid protein that forms part of the melanosome matrix, an important structural component of the mature melanosome [60–62]. Our candidate gene *Mlana* encodes a protein that interacts with Pmel and is also critical for melanosome matrix formation. We found that *Pmel* is downregulated in all Almond feather buds, and most severely in the two most depigmented types, light Almond and homozygous Almond (Fig. 5C). As described above, *Mlana* expression increased in dark Almond feathers but was similar to non-Almond in light Almond and homozygous Almond feather buds. These results are difficult to reconcile because these two genes are regulated by Mitf. Nevertheless, our results show that even the pigmented feathers in Almond birds show altered expression of pigmentation genes.

In summary, in homozygous Almond feather buds, the pigmentation production pathway is altered at an early stage of eumelanogenesis. In birds with one copy of the Almond allele (Z*^St^*Z*^+^* and Z*^St^*W) light feathers show downregulation of more eumelanin production genes than do dark feathers. Thus, phenotypically different Almond feathers have distinct pigmentation gene expression profiles.

### Other alleles at the *St* locus are copy number variants

Classical genetic studies point to multiple depigmentation alleles at the *St* locus [20, 29, 63, 64]. To determine if the Almond CNV is associated with these other alleles as well, we genotyped pigeons with other *St*-linked phenotypes and found significant increases in copy number in Qualmond (*St^Q^*; N=10, *p*=8.3e-06), Sandy (*St^Sa^*; N=3, *p*=3.2 e-02), Faded (*St^Fa^*; N=11, *p*=5.0e-07), and Chalky (*St^C^;* N=6, *p*=2.7e-04) pigeons compared to birds without *St*-linked phenotypes (Fig. 4, S4 Table). Another allele, Frosty (*St^fr^*), showed a trend of copy number increase that did not reach significance (N=6, *p*=1). Together, these results demonstrate that copy number increase is associated with a variety of depigmentation alleles at the *St* locus.

We next asked whether different *St* alleles share the same CNV breakpoints. We amplified and sequenced across the Almond CNV breakpoints in Qualmond (N=4), Sandy (N=2), Faded (N=2), and Chalky (N=4) pigeons and found that the breakpoints are identical in all phenotypes tested. Therefore, a single initial mutational event was probably followed by different degrees of expansion in different *St* alleles. Notably, the breakpoints of the 77-kb segment (ScoHet5_227: 5,181,467 and 5,259,256) are enriched for CT repeats. These repeat sites could facilitate non-allelic homologous recombination, which could have generated the *St* allelic series [65].

## DISCUSSION

### *Mlana* is a strong candidate gene for the Almond phenotype

We identified a CNV associated with plumage pigmentation variation and an eye defect in domestic pigeons. Different numbers of copies of this structural variant are associated with a series of depigmentation alleles at the same locus. In the feathers of Almond birds, the CNV is associated with changes in the expression of genes within its bounds.

One of these genes, *Mlana*, is a strong candidate for Almond due to its role in melanosome maturation. *Mlana* and *Pmel* are co-regulated by Mitf and their protein products physically interact with each other during the process of matrix formation in the melanosome [41, 66]. Notably, *Pmel* mutations cause pigmentation phenotypes in cattle, chicken, and mouse [67–70]. *Pmel* mutations in horse, dog, and zebrafish result in both pigmentation phenotypes and eye defects, similar to Almond pigeons [37, 71–75]. For example, the merle coat pattern in dogs is associated with a transposon insertion in an intron of the *PMEL* gene, resulting in a non-functional PMEL protein and a phenotype that is remarkably similar to the Almond phenotype in pigeons [37, 72]. Dogs homozygous for the *PMEL* mutation, much like homozygous Almond pigeons, are severely hypopigmented. Additionally, homozygous *PMEL* mutant dogs have various eye defects, such as increased intraocular pressure, ametropia, microphthalmia, and coloboma [76]. The observation that Pmel, which interacts directly with Mlana, is repeatedly connected to both pigmentation and eye defects makes *Mlana* a strong candidate for similar correlated phenotypes in Almond pigeons. Likewise, in humans and mice, mutations in melanosome genes (e.g., *Oca2*, *Slc45a2*, *Slc24a5*) produce both epidermal depigmentation and eye defects, thereby further demonstrating a shared developmental link between these structures [77–79].

The other full-length gene within the CNV, *Slc16a7*, does not have a known role in pigmentation. However, this gene is a member of a class of monocarboxylate transporters that are necessary to efficiently remove lactate from photoreceptor cells to prevent intracellular acidosis, and to maintain a high glycolysis rate and proper cellular metabolism [44–46, 80, 81]. We speculate that irregular expression of this gene could lead to cell death or dysfunction by causing toxic lactic acid concentrations or by preventing lactic acid transport to nearby cells. In regenerating Almond feathers, *Slc16a7* expression increases substantially (40-fold) relative to non-Almond feathers, raising the possibility that this gene is somehow involved in pigmentation. In short, changes in *Slc16a7* expression could drive components of the Almond phenotype in feathers, eyes, or perhaps both. However, given the linked pigment and eye phenotypes observed in *Pmel* mutants in other species, *Mlana* alone could be sufficient to induce both pigmentation and eye defects in Almond pigeons. Future work will explore these various possibilities.

### Gene expression is altered in Almond birds

In other organisms, copy number variation can result in gene expression changes in the same direction as the copy number change (i.e., the presence of more copies is correlated with higher expression) [82–84]. We observed a similar trend of higher expression of genes captured in the Almond-linked CNV (Fig. 5A). In contrast to this trend, however, *Mlana* showed an increase in expression in dark Almond feathers, but not in light Almond or homozygous Almond (unpigmented) feathers. *Mlana* is also the gene with the greatest copy number increase, with up to 14 copies in hemizygous Almond genomes and 28 copies in the homozygous Almond genome.

With the above observations of gene expression in mind, why might homozygous Almond birds lack *Mlana* expression in feather buds when they have 28 copies of the gene? One possibility is epigenetic silencing. High copy numbers in tandem arrays induce gene silencing in several organisms [85–89]. In fruit flies, for example, tandem arrays lead to variegated gene expression of the white eye gene [86]. This change in expression, in turn, leads to mosaic eye color, a scenario reminiscent of the color mosaicism in the feathers of Almond pigeons. In mouse, experimentally reducing the number of copies of *lacZ* in a tandem array causes an increase in gene expression, indicating that reducing copy number may relieve gene silencing [88]. Likewise, it is possible that somatic copy number decrease could relieve gene silencing and restore higher expression of *Mlana* in dark Almond feather buds.

Another potential explanation for the lack of *Mlana* expression in homozygous Almond feathers is cell death or immunity-mediated destruction of melanocytes. Overexpression of *Mlana* could have a toxic effect on cells, leading to cell death before melanocyte maturation. Similarly, in humans, overexpression of genes is often associated with disease [90–92], and in yeast, overexpression of genes can reduce growth rate [93]. Alternatively, Almond melanocytes might elicit an autoimmune response, similar to the destruction of melanocytes in human pigmentation disorders. MLANA is a dominant antigenic target for the T cell autoimmune response in human skin affected by vitiligo [94, 95], and perhaps the presentation of Mlana antigens in Almond pigeons elicits a response that depletes melanocytes in the developing feather buds. A potentially analogous autoimmune response depletes the melanocyte population and mimics vitiligo in Smyth line chickens [96].

If genes in the CNV are being randomly silenced in Almond pigeons, or cells with high expression are escaping cell death in a random manner, then we might expect to see high variance in gene expression among Almond feather samples. Consistent with this prediction, the variance in expression of *Mlana* in both dark and light Almond feather buds trends higher than in non-Almond samples (Fig. 5A). This variance might also explain the random pattern of pigmentation and de-pigmentation observed in the feathers of these birds. If each cell population is affected differently due to stochastic events resulting in differential expression, then random pigmentation patterns could be the outcome.

### CNVs as mechanisms for the rapid generation of new phenotypes

In addition to finding a CNV at the *St* locus in Almond birds, we found quantitative variation in copy number among other alleles at this locus. Variation at this CNV may have a quantitative effect on de-pigmentation, with the degree of copy number increase correlating with degree of depigmentation and eye defects. For example, pigeon breeders report that Sandy and Whiteout – two phenotypes with among the highest numbers of copies of the CNV (Fig. 4) – have associated eye defects similar to Almond (Tim Kvidera, personal communication) [29, 63]. Although we currently have a small sample size of other *St*-linked phenotypes, we see a trend that other alleles produce milder pigment phenotypes and have less CNV expansion than the Almond allele. Similar quantitative effects of CNVs occur in other organisms as well, including a correlation between comb size and copy number of *Sox5* intron 1 in chickens [97].

Pigeon breeders have reported that parents with one *St*-linked phenotype can produce offspring of another phenotype in the *St* series [29, 98]. Specifically, Faded, Qualmond, and Hickory pigeons have produced Almond offspring. These classical breeding studies suggest that allelic conversion can occur rapidly and, based on our finding of copy number variation among *St* alleles, may result from simple expansion or contraction of a CNV. In another striking similarity between Merle dogs and Almond pigeons, germline expansions or contractions of the *Merle* allele of *PMEL* result in a spectrum of coat pattern phenotypes that can differ between parents and offspring [72, 99]. Thus, unstable CNVs like the one we found at the *St* locus may provide a mechanism for extraordinarily rapid phenotypic diversification in pigeons and other organisms [100–103].

## MATERIALS & METHODS

### Animal husbandry

Animal husbandry and experimental procedures were performed in accordance with protocols approved by the University of Utah Institutional Animal Care and Use Committee (protocols 10-05007, 13-04012, and 16-03010).

### DNA sample collection and extraction

Blood samples were collected in Utah at local pigeon shows, at the homes of local pigeon breeders, and from pigeons in the Shapiro lab. Photos of each bird were taken upon sample collection for our records and for phenotype verification. Breeders outside of Utah were contacted by email to obtain feather samples. Breeders were sent feather collection packets and instructions, and feather samples were sent back to the University of Utah along with detailed phenotypic information and genetic relatedness. DNA was then extracted from blood, as previously described [4]. DNA from feathers was extracted using the user developed protocol for Purification of total DNA from nails, hair, or feathers using the DNeasy Blood & Tissue Kit (Qiagen Sciences, Germantown, MD).

### Genomic analyses

BAM files from a panel of previously resequenced birds were combined with BAM files derived from new sequences from 11 Almond females and 16 non-Almond birds aligned to the Cliv_2.1 genome assembly [104] (new sequence accessions: SRA SRP176668, accessions SRR8420387-SRR8420407 and SRR9003406-SRR9003411; BAM files created as described previously [6]). SNVs and small indels were called using the Genome Analysis Toolkit (Unified Genotyper and LeftAlign and TrimVariants functions, default settings [105]). Variants were filtered as described previously [38] and the subsequent variant call format (VCF) file was used for downstream analyses.

Whole genomes of 12 Almond and 96 non-Almond birds were tested for allele frequency differentiation using pFst (VCFLIB software library, https://github.com/vcflib; see S1 Table for sample information) [38]. For analysis of fixed coding changes, VAAST 2.0 [39] was used to conduct an association test and to search for putative disease-causing genetic variants common to all Almond individuals but absent from non-Almonds. Annotated variants from affected individuals were merged by simple union into a target file. The background file included variants from 66 non-Almond birds, while the target file contained variants from the 12 Almond birds. This VAAST analysis revealed that there were no fixed genetic variants among the Almond individuals that were absent in the background dataset.

### CNV breakpoint identification and read-depth analysis

Read depth in the CNV-containing region was analyzed in 12 Almond and 118 non-Almond resequenced whole genomes. Scaffold ScoHet5_227 gdepth files were generated using VCFtools [108]. Read depth was normalized using a region (scaffold ScoHet5_227: 1-5,000,000) that did not show an increase in sequencing coverage in Almond genomes.

To determine the CNV breakpoints, we first identified the region of increased sequencing coverage in Almond genomes using the depth function in VCFtools [108]. Next, we examined BAM files of Almond genomes in IGV [110] in the region of coverage increase, and identified locations at which reads were consistently split (did not map contiguously). These locations were the putative breakpoints. We then designed PCR primers that amplify 1-kb products spanning the putative breakpoints (see S5 Table for primer sequences). Finally, we used PCR to amplify across the putative breakpoints. PCR products were purified and sequenced, and aligned to the pigeon genome assembly using Blast+ version 2.7.1 [109]. The CNV breakpoint primers (see Fig. 3B) successfully amplified products in 40 of 43 Almond pigeons tested.

### Fusion gene analysis

The putative mRNA sequence of the *Ermp1/Kiaa2026* fusion gene was determined by concatenating the mRNA sequence of the exons on one side of the outer breakpoint with the exons that map to the outer breakpoint. The fusion of these exons was confirmed using exon spanning primers and qPCR (See S5 Table for primer sequences). The putative mRNA sequence was translated, and then analyzed for domains using HMMER searches in SMART (Simple Modular Architecture Research Tool) [48]. We searched for domains in the SMART database, and also searched for outlier homologs, PFAM domains, signal peptides, and internal repeats.

### Taqman assay for copy number estimates

Copy number variation was estimated using a custom Taqman Copy Number Assay targeted to the *Mlana* region (MLANA_CCWR201) for 150 Almond, 9 Qualmond, 3 Sandy, 14 Faded, and 6 Chalky, 5 Frosty, and 56 individuals without *St*-linked phenotypes. Following DNA extraction, samples were diluted to 5 ng/uL and run in quadruplicate according to manufacturer’s protocol. Copy number was determined using CopyCaller Software v2.1 (ThermoFisher Scientific, Waltham, MA). An intron in *RNaseP* was used for normalization of copy number.

### RNA isolation and cDNA synthesis

To assay gene expression, secondary covert wing feathers were plucked to stimulate regeneration and allowed to regenerate for 9 days (see S2 Table for sample details). Nine-day regenerating feather buds were plucked, then the proximal 5 mm was cut and stored in RNA later at 4°C overnight. Feather buds were then dissected and collar cells removed, and stored at −80°C until RNA isolation. RNA was then isolated and reverse transcribed to cDNA as described previously [4].

### qRT-PCR analysis

cDNA was amplified using intron-spanning primers for the appropriate targets using a CFX96 qPCR instrument and iTaq Universal Sybr Green Supermix (Bio-Rad, Hercules, CA) (S5 Table). Samples were run in duplicate and normalized to β-actin (see S3 Table for raw results). Results were compared in R [111] using ANOVA, followed by a Tukey post hoc test to determine differences between phenotypic groups. Differences were considered statistically significant if *p* < 0.05. Primers used for each gene are included in S5 Table.

## ACKNOWLEDGEMENTS

We thank past and present members of the Shapiro lab for assistance with sample collection and processing; members of the Utah Pigeon Club and National Pigeon Association for sample contributions; and Ken Davis and Tim Kvidera for critical discussions and advice. We thank Tim Kvidera for photographs of Whiteout, Sandy, Frosty, Faded, and Chalky pigeons in Figure 4. We thank Anna Vickrey, Max Sidesinger, Elena Boer, Emily Maclary, Sara Young, and Robert Greenhalgh, for technical assistance and advice. This work was supported by the National Science Foundation (CAREER DEB-1149160 to M.D.S.; GRF 1256065 to R.B.) and the National Institutes of Health (R01GM115996 and R35GM131787 to M.D.S., R01GM104390 to M.Y., fellowship T32GM007464 to Z.K.). The funders had no role in study design, data collection and analysis, decision to publish, or preparation of the manuscript. We acknowledge a computer time allocation from the Center for High Performance Computing at the University of Utah.

## SUPPORTING INFORMATION LEGEND

**Supporting_Information.xlsx.** S1-S5 Tables. Individual tables are separate worksheets within the file and contain the following information: S1, NCBI SRA submission numbers and breed information for birds used for genomic analysis in this study. S2, Sample sizes and Identifiers of birds included in each phenotypic category for qPCR analysis. S3, Raw qRT-PCR results for Figure 4. S4, Copy number results from Taqman assay of *Mlana* region. S5, Primer Sequences Used in This Study. S2:

